# Approach to insect wing shape and deformation field measurement

**DOI:** 10.1101/119230

**Authors:** Duo Yin, Zhen Wei, Zeyu Wang

## Abstract

**Summary Statement:** A fine shape and deformation field measurement of insect wing is achieved by a self-developed setup. This measurement could foster investigation of insect wing stiffness distribution.

**Abstract:** For measuring the shape and deformation of insect wing, a scanning setup adopting line laser and coaxial LED light is developed. Wing shape can be directly acquired from the line laser images by triangulation. Yet the wing deformation field can also be obtained by a self-devised algorithm that processes the images from line laser and coaxial LED simultaneously. During the experiment, three wing samples from termite and mosquito under concentrated force are scanned. The venation and corrugation could be significantly identified from shape measurement result. The deformation field is sufficiently accurate to demonstrate its variation from wing base to tip. The load conditions in experiments are also be discussed. For softer wings, local deformation is apparent if pinhead is employed to impose force. The similarity analysis is better than 5% deformation ratio as a static criterion, if the wing is simplified as a cantilever beam. The setup is proved to be effective and versatile. The shape and deformation fields would give enough details for the measurement of wing stiffness distribution.

## Introduction

Insect wings undergo complex and even large deformation when propelling insects into the air(Wootton, 1990). The insect wing deformation in flight affects flight control and lift generation on a big scale. The understanding of wing deformation in flight could dramatically foster the design of flexible wing for flapping wing micro air vehicle (FWMAV) in engineering circles. The FWMAV has considerable advantages than traditional micro air vehicle in size, survivability and controllability. Several FWMAVs inspired by insects prove to be promising, including the Micromechanical Flying Insect (Fearing, 2004; Yan et al., 2001), DelFly (de Croon et al., 2009; de Croon et al., 2012; Lentink et al., 2007) and Harvard Microrobotic Fly (Ma et al., 2013). Remarkable progress of the FMAV has been achieved on energetics (Karpelson et al., 2010), actuation (Wood et al., 2005)and aerodynamics (Deng et al., 2014; Nakata et al., 2011; Wilkins and Knowles, 2009). Yet the flight performance of FWMAV can hardly match with that of the insects.

The wing deformation in flight is determined by both load that wing carries and the wing stiffness. The load could be roughly categorized as aerodynamic force and inertial-elastic force(Daniel and Combes, 2002). Both these two forces have proved to be crucial factors for the instantaneous shape of the wing(Ellington, 1984; Ennos, 1988; Ennos, 1989; Wilkin and Williams, 1993; Zanker and Gotz, 1990). The wing stiffness distribution is complicated because the wing is a passive structure mainly composed of veins and membranes (Wootton, 1992) and with no internal muscles inside (Mengesha et al., 2011). Although several researches focusing on measuring flexural stiffness distribution of insect wing have been done (Combes and Daniel, 2003; Ganguli et al., 2010; Lehmann et al., 2011; Mengesha et al., 2011), the wing is simplified as a one-dimensional beam in these researches, thus the spatial flexural stiffness distribution was not measured precisely. To acquire fine spatial flexural stiffness distribution requires an accurate deformation measurement first. Meanwhile, an exact shape measurement of the wing is the first step in the quest of a precise deformation measurement.

Several accurate methods have been adopted in measuring insect wing shape, including interferometry, micro CT and laser triangulation. Based on laser interferometry method, a three-dimensional shape measurement system was designed to measure surface roughness of mosquito *Culicidae* wing (Sudo et al., 2005; Sudo et al., 2000). The measurement accuracy could reach submicron. The dragonfly *Anisoptera* wing shape was investigated via micro CT (Jongerius and Lentink, 2010). The scanning resolution is 7.2 μm. Thickness and shape measurement (TSM) method was devised to measure thickness and shape of dragonfly *Anisoptera* wing simultaneously (Zeng et al., 1996). The TSM method is composed of heterodyne interferometry and laser triangulation. The error of their measurement is within 1.8%. Although these researches could obtain fine shape information, none of them has given a point to point matching deformation field.

In the current paper, the laser stripe triangulation and image matching are employed to measure the shape and deformation of the insect wing under concentrated force. A setup, based on triangulation, is designed and fabricated to achieve the measurement. By light stripe center extraction and image matching, the wing shape and deformation field could be obtained.

## Materials and Methods

### Study specimens

The tested wing specimens are wings from formosan subterranean termite *Coptotermes formosanus* and two kinds of mosquito *Culicidae.* These two *Culicidae* are not identified the exact species due to limitation of sampling. In this article, the two *Culicidae* are written as *Culicidae 1* and *Culicidae 2* in short. All these insects are collected in local garden in summer. They are common flying insects in the vicinity. These wings are chosen for the significant difference in size or venation.

### Triangulation and setup

Laser strip triangulation and grayscale square centroid method are used in the wing shape measurement. Triangulation is based on trigonometry. It is highly flexible, it can measure large object like the seafloor roughness (Wang and Tang, 2009) and small object like nano-composite ceramic coatings (Portinha et al., 2003). In this research, laser stripe triangulation is adopted for its robustness and versatility.

Based on triangulation, a setup has been designed for the measurement. The setup is mainly composed of camera, lens, coaxial LED, line laser, linear guide for samples, motion stage and microforce sensing probe. The coaxial LED light and line laser are all in 405nm wavelength. The coaxial LED illuminates wing sample at right angle, while the line laser lightens it with an incidence angle about 45^o^. The linear guide could drive the whole wing sample pass through the measurement region step by step. A microforce sensing probe on the adjustable motion stage could impose a precise concentrated force to the wing sample. Fig. 1 shows the measurement setup.

**Fig. 1.**
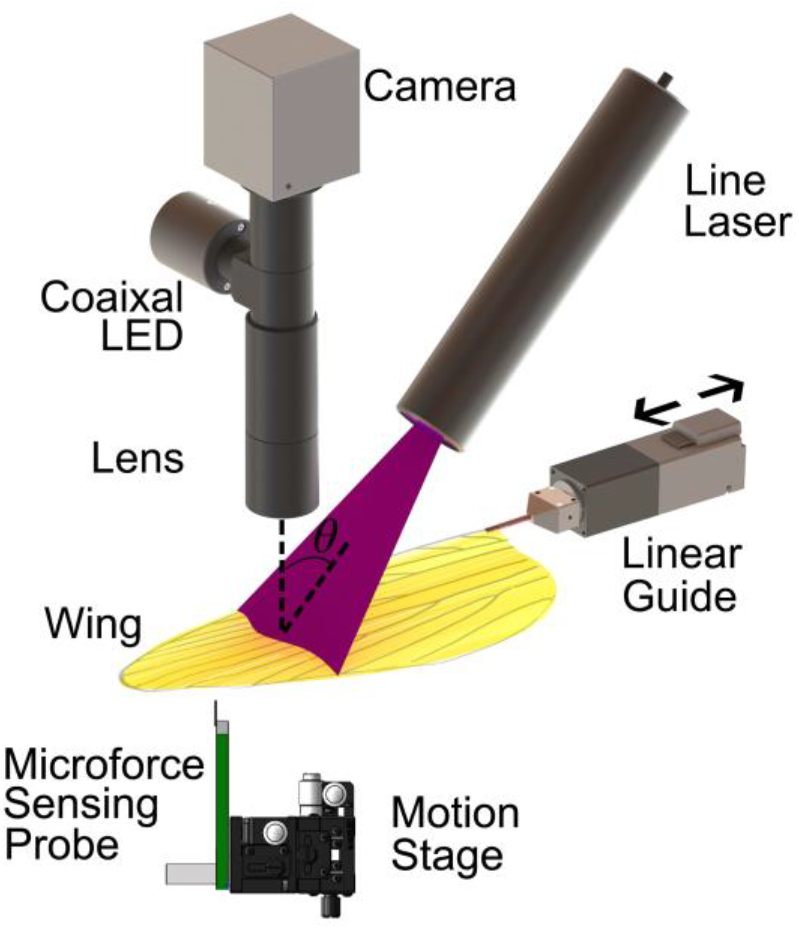
Measurement setup. Main components of the setup are shown, including camera, line laser, coaxial LED, lens, linear guide, motion stage and mocroforce sensing probe.

During the experiment, the wing fixed on the setup would be scanned twice. In the first scanning, the wing sample, receiving no concentrated force, would be scanned step by step under the illumination of coaxial LED and line laser alternately till the end of the whole wing. Before the second scanning, the microforce sensing probe would carefully impose a very small concentrate force on the wing sample surface. As a result, the wing would have a very small deformation. Then the wing would be scanned again like the first scanning.

The setup should be accurately calibrated before measurement. The primary calibration parameters include physical sizes per pixel in each direction, laser stripe width and its incidence angle. A machine vision chessboard, a low reflection flat plate and a self-developed 3D step-shaped plate are used to determine the calibration parameters above. Physical sizes per pixel in each direction and laser incidence angle can be calculated by scanning the chessboard and 3D step-shaped plate. The laser stripe width can be obtained by measuring the laser strip on the low reflection flat plate.

### Shape measurement

While the laser illuminates the wing, the image with an irregular light strip could be recorded. By processing this image, three-dimensional coordinates of a series points on the wing can be calculated. Therefore, the cloud points of the whole wing sample can be built if enough images for different wing parts are collected and processed. In the light strip processing, the grayscale centroid method is applied because it could provide a high quality result. Assuming that (*cX_j_, cY_j_*) is the strip center position in pixel at the row *j* on the image, it could be calculated

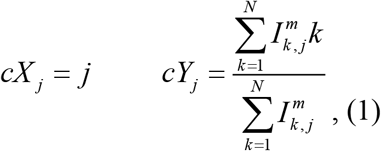

In Eqn.1, *I_k,j_* is the grayscale of the pixel on image position (*k,j*). *m* is the grayscale power weight factor and its value is 2 here. N is the row size of image. If (*cX_j_, cY_j_*) of each row *j* is ready, its three-dimensional coordinates can be derived from the calibration results. The generation of NURBS surface from cloud points is a standardized procedure and would not be described here.

### Deformation measurement

The key of deformation measurement is to find the matching positions of the points before and after deformation. A combination of triangulation and image matching method is employed here. Triangulation could provide coordinate of each point and image matching could find its corresponding matching point after deformation.

Only triangulation cannot provide the correct matching relationship during the deformation measurement. The points lighten by laser before deformation will not be lightened again on the same step after deformation. In fact, the real matching point would be lightened by laser on another step after deformation, so its position cannot be obtained only by triangulation if the matching step is uncertain. The images illuminated by LED are gray images which can give the wing surface details around laser strip on the same step, thus the real matching point can be found by using SIFT algorithm to analyze these gray images.

Fig. 2 illustrates the positions of matching points. *A_0_* is a point lightened by laser in step *i* before deformation. *A_2_* is the point lightened by laser in the same step after deformation. According to analysis above, *A_2_* is not the matching point of *A_0_.* The matching points for gray images are both *A_1_* in step *i* and *A^*^* in another step *j*. *A_1_* is the final right matching point for deformation calculation but it is not lightened by laser in step *i.* Fortunately, the position of *A^*^* could be calculated by triangulation and from *A^*^* to *A_1_* only has a calculable rigid displacement. Finally the position of matching point in same step *A_1_* could be easily derived from *A^*^*.

**Fig. 2.**
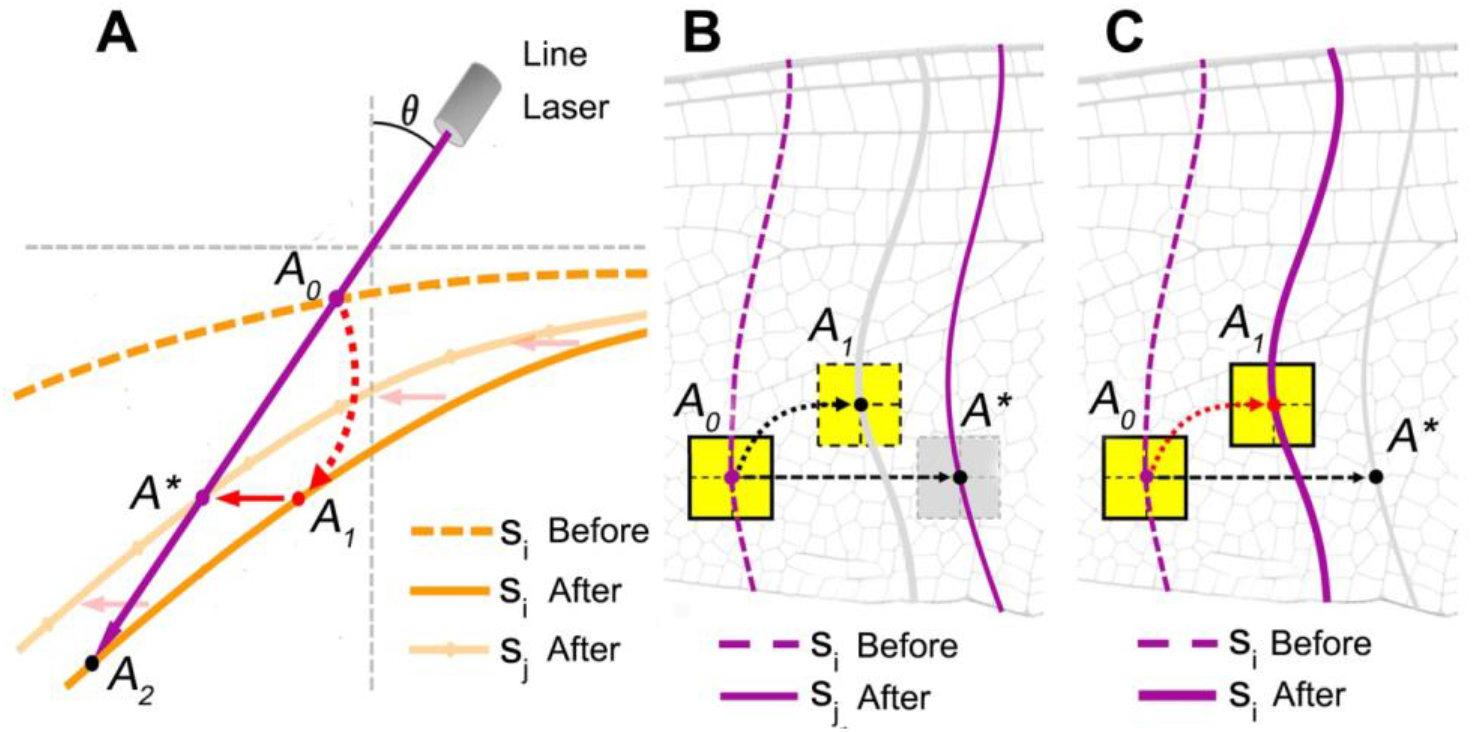
Matching point acquisition. (A) Relationship of matching points. The dotted and solid orange lines are wing shape in step *i* before and after deformation. Solid light orange line is wing shape in another step *j* after deformation. *A_0_* and matching point *A\** are on dotted orange line and solid light orange line re spectively. *A_2_* and matching point *A_1_* are on solid orange line. (B) Position deter mination of matching points *A_0_* and *A*.* Dotted purple line is laser stripe in step *i* before deformation. The solid purple line is laser stripe in another step *j* aft er deformation. The blocks in yellow and gray are gray image blocks centered i n matching points. (C) Position determination of matching point *A_0_* and *A_1_.* The dotted and solid purple lines are laser stripe in step *i* before and after deforma tion. The yellow blocks are gray image blocks centered in matching points.

## Results

### Setup

The main components of the setup are camera, lens, linear guide and microforce sensing probe. The adopted camera, MVUB500M, is an industrial camera. MML08-ST170D is employed as the lens. KR20, by THK Co., Ltd., is chosen as the linear guide for its high accuracy. FT-S1000 is used as the microforce sensing probe.

### Calibration

Calibrations for physical sizes per pixel and light stripe width are similar to other triangulation calibration methods. To determine the incidence angle, a step-shaped template is designed and fabricated. The template is a slim bar with four steps. The step height is 0.05 mm. In the experiment, when the laser scans the template, there emerges a convex on laser image. The convex height represents displacement caused by step height. The incidence angle could be calculated from step height and convex height. Table 1 shows the calibration result.

**Table 1.** Calibration result. Light stripe width, physical sizes per pixel and incidence angle are calibrated.

| Light stripe width (μm) | Physical sizes per pixel (mm) | Incidence angle ( ° ) |
| --- | --- | --- |
| 25.6±11 | 3.53e<sup>-3</sup> | 43.86 |

### Shape measurement

Wings from formosan subterranean termite *Coptotermes formosanus* and two kinds of mosquito *Culicidae* are chosen as the wing samples. These wing samples are different in shape and size. The wing lengths of *Coptotermes formosanus, Culicidae 1, Culicidae 2* are about 13 mm, 11 mm and 3 mm respectively. Wings from dead body are scanned for their stability of mechanical property. Fig. 3A-C is the wing sample photograph.

**Fig. 3.**
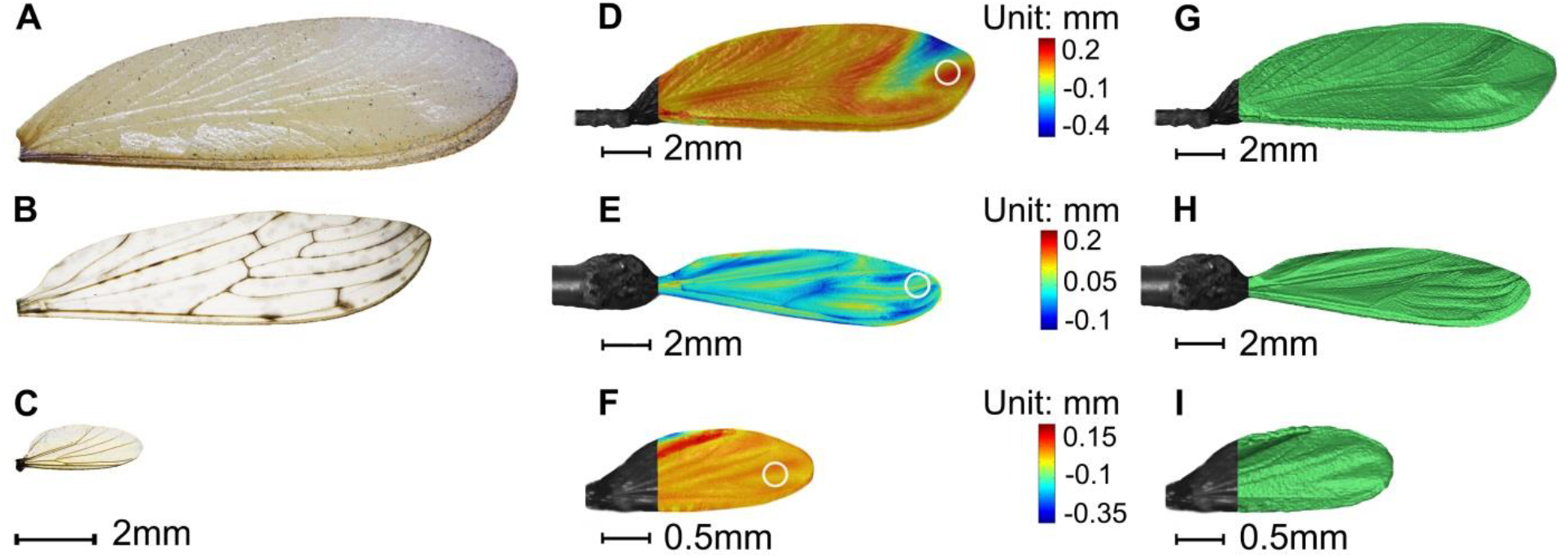
Real wing shape and measured wing shape. (A-C) Photographs of *Coptotermes formosanus, Culicidae 1* and *2* wings. These insect samples are obtained in the local garden. (D-F) Shape contours of *Coptotermes formosanus, Culicidae 1* and *2* wings. The contours are superimposed on the corresponding gray images. The white ring represents the probe. Its size and position reflect probe size and load position. (G-I) NURBS surfaces of *Coptotermes formosanus*, *Culicidae 1* and *2* wings. The NURBS surfaces are also superimposed on the gray images.

During the experiment, the wing with a rod is fixed on the sample holder of scanning system, then it is scanned twice. In the first scanning, both LED and line laser are used to scan the samples. The scanning step length is 0.01 mm. Before the second time scanning, the microforce sensing probe imposes a concentrated force to wing tip area. The white rings in Fig. 3D-F represent force position and probe size. The force position is near leading edge tip for *Culicidae 2* wing and near to trailing edge for *Coptotermes formosanus* and *Culicidae 1* wings. The force is not measured but the wing deformation due to this force is limited to a small scale. After two times scanning, the wing shape and its deformation can be obtained by using methods above. Sometimes, the wing cannot be perfectly scanned due to the reflecting characteristics of the wing. For *Coptotermes formosanus* and *Culicidae 2* wing samples here, wing base reflection is too strong to extract the accurate center, thus the wing base is not shown in the result.

Fig. 3G-I shows the wing shape contour and NURBS surface from the first scanning. Significant vein patterns and shape characteristics could be observed. The *Coptotermes formosanus* wing tip is sunken near trailing edge. The wing of *Culicidae 1* is flat. The trailing edge of *Culicidae 2* wing is arched. The membrane would undergo wrinkling during natural dehydration, but the spatial distribution of vein is clear enough in the result.

### Deformation measurement

Fig. 4-6 summarizes the deformation distribution for all wing samples. Both the deformation contour and deformation profiles along spanwise and chordwise are shown. The origin of the coordinate system locates near wing base. The x axis is along spanwise from root to tip while y axis points from leading to trailing edge. The deformation is filtered by wavelet filter. The matching point correlation coefficient distribution for its displacement calculation is also filtered and outlined. It is a useful value to identify the accuracy of deformation measurement. The correlation coefficient is high enough (few less than 0.75) for all wing samples, so the deformation measurement is fine. The root and tip are also dropped in correlation and deformation calculation for the same reflection influence.

The deformation fields indicate that the wing can be considered as a cantilever beam with bending and torsion under the concentrated force. According to the results, the small deformation condition is still valid, and the samples are in elastic status, but the deformation is more complex than a one dimensional simple beam.

For *Coptotermes formosanus* wing in Fig. 4, the maximum deformation value can be found near the probe contact area, the amount of deformation is about 0.30 mm, thus the deformation ratio is about 2.3%. Two additional high deformation zones near trailing edge can also be specified, where the deformation values are over 0.11 mm. The deformations on rest areas are mostly under 0.10 mm. The zones with deformation over 0.15 mm are all concentrated in the region of probe position.

**Fig. 4.**
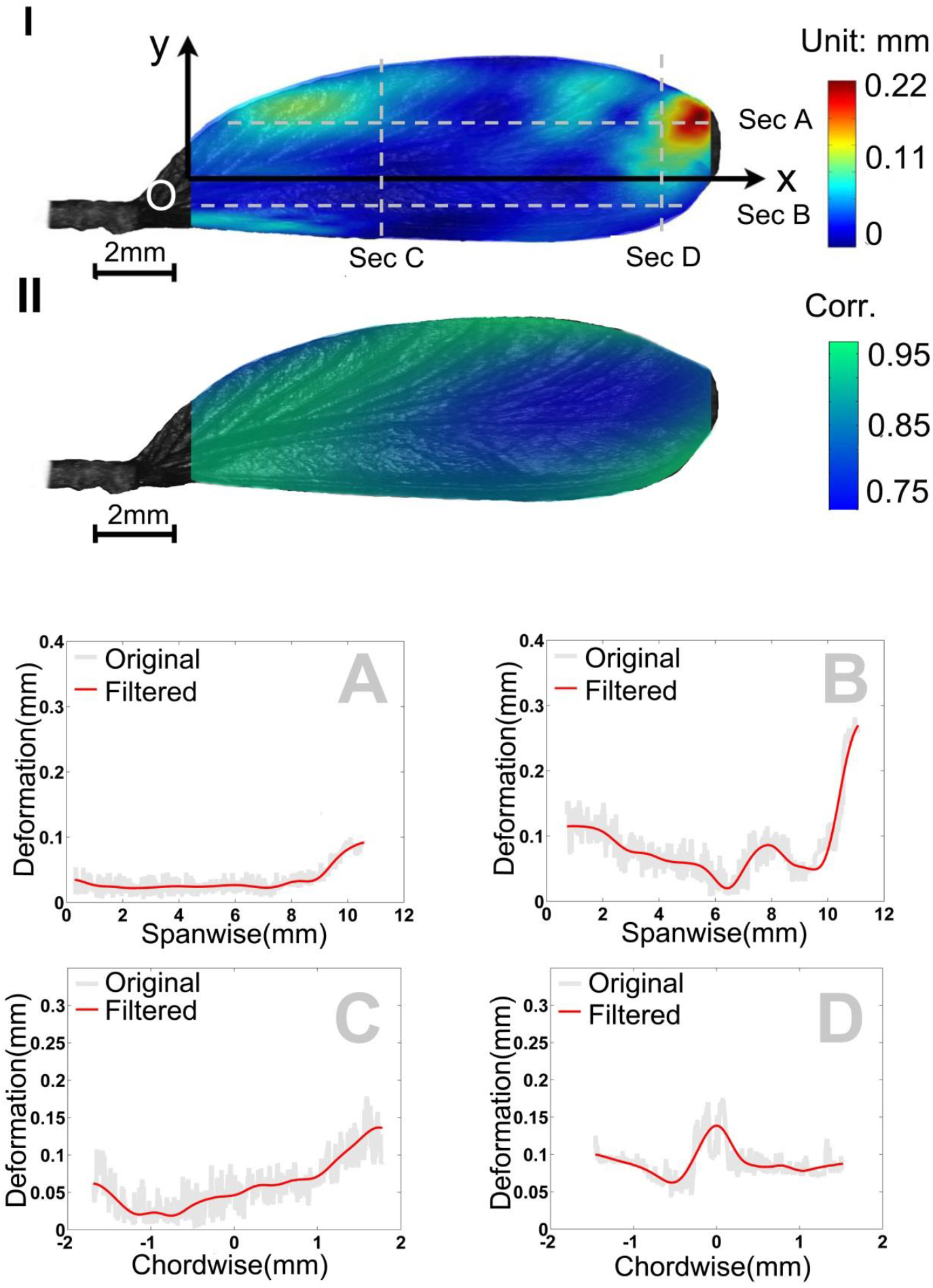
*Coptotermes formosanus* wing deformation. (I) Deformation contour. The deformation value is wavelet filtered. (II) Correlation coefficient contour. The coefficient value is wavelet filtered. (A-D) Deformation profiles along sections A-D in Fig. 4I. The gray and red lines represent original deformation value and wavelet filtered one respectively.

Fig. 4A shows the deformation near leading edge. Before 60% spanwise length on Section A, the amount of deformation is just 0.05 mm and almost no increase. Near the wing tip on Section A, the deformation mildly increases to 0.1 mm. While the deformation near trailing edge (Fig. 4B) varies intensively. From wing base to tip along the Section B, the deformation value goes down to zero from 0.11 mm and then increases to 0.28 mm. The deformation near wing base (Fig. 4C) changes gradually, and has peak value at about 0.13 mm near trailing edge. The deformation near load position (Fig. 4D) emerges an obvious at peak about 0.15 mm in the middle because it is nearly cross over the probe zone.

For *Culicidae 1* wing in Fig. 5, the spanwise deformation trend increases gradually. There is no significant different between spanwise deformation near leading edge (Section A) and that near trailing edge (Section B). As chordwise deformation near wing base (Section C) is small and flat, torsion contributes little to the whole deformation.

**Fig. 5.**
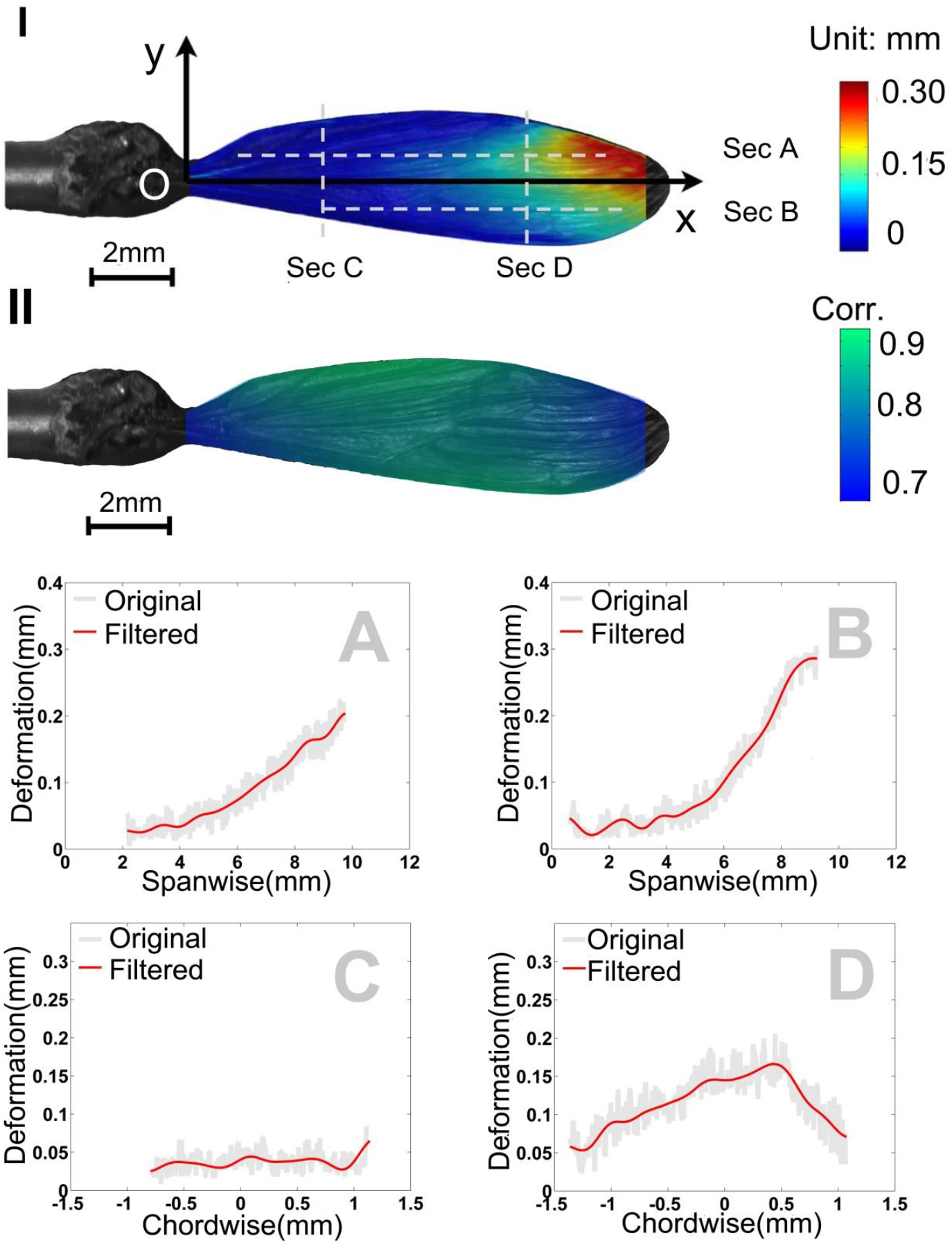
*Culicidae 1* wing deformation. (I) Deformation contour. The deformation value is wavelet filtered. (II) Correlation coefficient contour. The coefficient value is wavelet filtered. (A-D) Deformation profiles along sections A-D in Fig. 5I. The gray and red lines represent original deformation value and wavelet filtered one respectively.

The maximum deformation value of *Culicidae 1* wing is about 0.30 mm, so the deformation ratio is about 2.7%. The peak deformation near leading edge (Fig. 5A) is around 0.22 mm, with no flat or decrease from wing base to tip. Correspondingly, the peak is about 0.29 mm near trailing edge (Fig. 5B). The deformation near wing base (Fig. 5C) has no obvious peak and the value is below 0.05 mm, while on profile near load position (Fig. 5D), there is a small peak value at about 0.16 mm. The *Culicidae 1* wing deformation is more regular than *Coptotermes formosanus* wing despite they have similar load condition.

As shown in Fig. 6, the maximum deformation value of *Culicidae 2* wing is about 0.1 mm, the deformation ratio is still just about 3.3% even wing size is only 3.0 mm. Similar to the *Coptotermes formosanus* wing deformation, the zone with greater deformation is in the same region of the probe position. Different with *Coptotermes formosanus* wing deformation, *Culicidae 2* wing only has one high deformation zone.

**Fig. 6.**
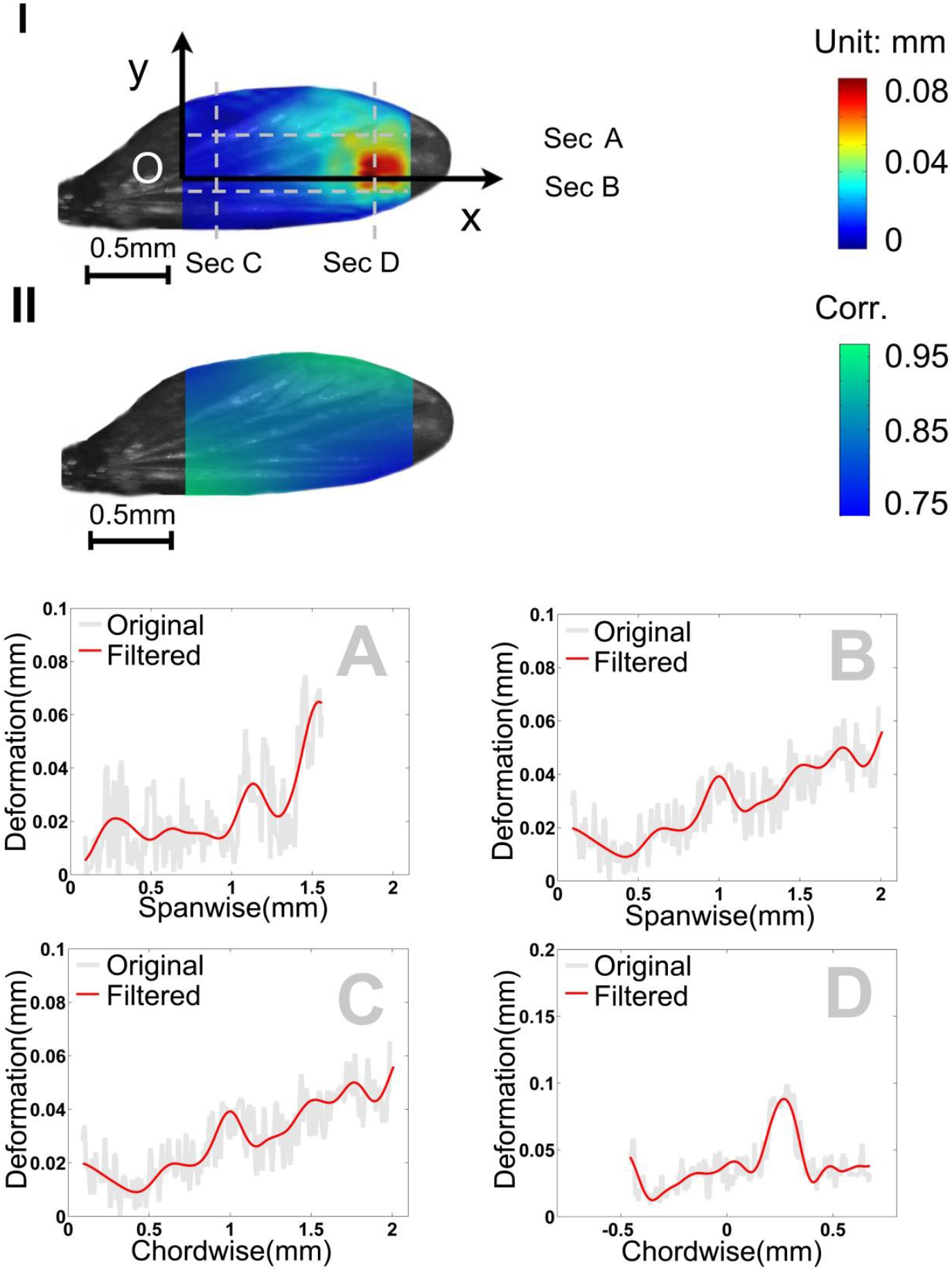
*Culicidae 2* wing deformation. (I) Deformation contour. The deformation value is wavelet filtered. (II) Correlation coefficient contour. The coefficient value is wavelet filtered. (A-D) Deformation profile along sections A-D in Fig. 6I. The gray and red lines represent original deformation value and wavelet filtered one respectively.

Fig. 6A-D shows the deformation profiles of *Culicidae 2* along sections A-D. The peak deformation near leading edge (Section A) is around 0.07 mm. The deformation trend resembles section A of *Culicidae 1* wing, but with more volatility from wing base to tip. The deformation variation near trailing edge (Section B) is similar with that near leading edge (Section A), but the peak value is 50% lower because section A is closer to the probe zone. The deformation near wing base (Section C) is similar to the section C of *Culicidae 1* wing as well, and its peak value is about 0.05 mm at trailing edge point. The deformation near load position (Section D) has a peak value at about 0.1 mm in the middle because it locates in the probe zone center. From the section comparison, it is clear that *Culicidae 2* wing deformation is more like that of *Culicidae 1* wing although there is concentrated deformation near the probe zone.

## Discussion

Insect wing shape and deformation measurement method presented in this article has been proved. The scanning can provide fine shape and deformation results for wings with from spanwise length (3 mm) to large one (13 mm). The uneven shape and vein pattern could be observed clearly. The wing deformation could be found precisely. It has been confirmed that the illumination by 405nm wavelength light is suitable for insect wing scanning, despite the images near the wing base is still imperfect. In fact, not like single or multiple point measurement before, it is a practical and convenient way to obtain deformation field. Therefore, it provides us a possibility to analyze the stiffness distribution more precisely by using methods like finite element analysis to matching the deformation field result.

By comparing the wing deformation results of *Coptotermes formosanus* and *Culicidae 1,* it could be found that the load condition should be carefully chosen. There are two typical loading methods for insect wing deformation and stiffness research, one is using a pinhead to impose a concentrated force on the wing surface(Combes and Daniel, 2003), and the other is using a knife edge to impose line load and restrict torsion in the wing bending(Ganguli et al., 2010). *Coptotermes formosanus* wing is relatively softer than *Culicidae 1* wing due to its material and structure, so it has a large deformation zone at the position of the probe. The *Culicidae 1* wing sample has almost the same spanwise length, aspect ratio, and it is also under the same load condition (0.3 mm maximum deformation), but its deformation is not concentrated. Thus if softer wing has been investigated, pinhead loading may results in local deformation. Correspondingly, for more rigid wing like *Culicidae 1* wing, both pinhead and knife edge loading may not induce any local deformation when small deformation assumption is satisfied. The deformation field is a valuable tool for checking whether the load condition is suitable or not.

By comparing the wing deformation measurement results of *Culicidae 1 and Culicidae 2,* it could also be found that the load condition should be carefully chosen. The deformation changing trend and vein structure of *Culicidae 1* wing are basically similar with those of *Culicidae 2* wing sample. The mainly difference is that *Culicidae 2* wing sample shows concentrated deformation. The difference is determined primarily by the load condition. Here a model of one dimensional cantilever beam with concentrated force at free end and similarity analysis are employed to explain this phenomenon. First of all, an equation of slope for cantilever beam can be given as below:

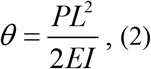

Where, P is the concentrated force, L is the beam length, I is the moment of inertia, and E is Young’s Modulus. If the wings of *Culicidae 1* and *Culicidae 2* have similar deformation distribution, they should have same slope value, so that:

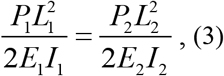

In Eqn. 3, the variables with suffix ‘1’ refer to *Culicidae 1* wing and the variables with suffix‘2’ refer to *Culicidae 2 wing.* The maximum deflection of such a cantilever beam can also be directly given as below:

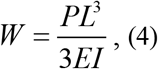

The maximum deflection of both *Culicidae 1* wing and *Culicidae 2* wing can be solved by Eqn. 4. The maximum deflection ratio between these two wings is derived as:

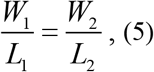

The meaning of Eqn. 5 is quite simple: only if wings have the same deformation ratio, they would have similar deformation distribution. In this case, the deformation ratio of *Culicidae 2* wing is a little bigger than that of *Culicidae 1* wing, so it may lead to different distribution. The discussion here is under the assumption that both geometry and material property are similar. Similarity analysis can be applied to wings of *Culicidae 1* and *Culicidae 2,* as they are both in *Diptera* order. The *Coptotermes formosanus* wing has similar deformation ratio as the *Culicidae 1* wing, but it doesn’t fulfil the similarity requirement because it is in the order of *Isoptera.* Normally, the deformation or stiffness experiments of insect wings use 5% small deformation assumption as the criterion. In fact, it is not enough if a regular deformation is required to satisfy the beam bending model. Similarity analysis should be considered when the load condition is changing.

## Acknowledgements

The author would express gratitude to Chengwu Wang and Changqiu Zhou in animal flight group of Chongqing University for their suggestion on image processing methods.

## Author contributions

Zhen Wei and Duo Yin devised the measurement setup. Zeyu Wang and Duo Yin obtained the insect samples from local garden. Duo Yin and Zeyu Wang performed the experiment and analyzed the experiment results. Duo Yin and Zhen Wei wrote the manuscript.

## Data availability

None

## References

Combes, S. A. and Daniel, T. L. (2003). Flexural stiffness in insect wings I. Scaling and the influence of wing venation. Journal of Experimental Biology 206, 2979-2987.

Daniel, T. L. and Combes, S. A. (2002). Flexible wings and fins: Bending by inertial or fluid-dynamic forces? Integrative and Comparative Biology 42, 1044-1049.

de Croon, G., de Clercq, K. M. E., Ruijsink, R., Remes, B. and de Wagter, C. (2009). Design, aerodynamics, and vision-based control of the DelFly. International Journal of Micro Air Vehicles 1, 71-97.

de Croon, G., Groen, M.A., De Wagter, C., Remes, B., Ruijsink, R. and van Oudheusden, B. W. (2012). Design, aerodynamics and autonomy of the DelFly. Bioinspiration & Biomimetics 7, 16.

Deng, S. H., Percin, M., van Oudheusden, B., Remes, B. and Bijl, H. (2014). Experimental Investigation on the Aerodynamics of a Bio-inspired Flexible Flapping Wing Micro Air Vehicle. International Journal of Micro Air Vehicles 6, 105-115.

Ellington, C. P. (1984). The aerodynamics of hovering insect flight .6. lift and power requirements. Philosophical Transactions of the Royal Society of London Series B-Biological Sciences 305, 145-181.

Ennos, A. R. (1988). The inertial cause of wing rotation in diptera. Journal of Experimental Biology 140, 161-169.

Ennos, A. R. (1989). Inertial and aerodynamic torques on the wings of diptera in flight. Journal of Experimental Biology 142, 87-95.

Fearing, R. S. (2004). Biological inspiration for micro flight: The micromechanical flying insect. Abstracts of Papers of the American Chemical Society 227, U525-U525.

Ganguli, R., Gorb, S., Lehmann, F.O. and Mukherjee, S. (2010). An Experimental and Numerical Study of Calliphora Wing Structure. Experimental Mechanics 50, 1183-1197.

Jongerius, S. R. and Lentink, D. (2010). Structural Analysis of a Dragonfly Wing. Experimental Mechanics 50, 1323-1334.

Karpelson, M., Whitney, J.P., Wei, G.Y., Wood, R.J. and Ieee. (2010). Energetics of Flapping-Wing Robotic Insects: Towards Autonomous Hovering Flight. In Ieee/Rsj 2010 International Conference on Intelligent Robots and Systems. New York: Ieee.

Lehmann, F. O., Gorb, S., Nasir, N. and Schutzner, P. (2011). Elastic deformation and energy loss of flapping fly wings. Journal of Experimental Biology 214, 2949-2961.

Lentink, D., Bradshaw, N. and Jongerius, S. R. (2007). Novel micro aircraft inspired by insect flight. Comparative Biochemistry and Physiology a-Molecular & Integrative Physiology 146, S133-S134.

Ma, K. Y., Chirarattananon, P., Fuller, S.B. and Wood, R. J. (2013). Controlled Flight of a Biologically Inspired, Insect-Scale Robot. Science 340, 603-607.

Mengesha, T. E., Vallance, R.R. and Mittal, R. (2011). Stiffness of desiccating insect wings. Bioinspiration & Biomimetics 6, 8.

Nakata, T., Liu, H., Tanaka, Y., Nishihashi, N., Wang, X. and Sato, A. (2011). Aerodynamics of a bio-inspired flexible flapping-wing micro air vehicle. Bioinspiration & Biomimetics 6, 11.

Portinha, A., Teixeira, V., Monteiro, A., Costa, M.F., Lima, N., Martins, J. and Martinez, D. (2003). Surface analysis of nanocomposite ceramic coatings. Surface and Interface Analysis 35, 723-728.

Sudo, S., Tsuyuki, K. and Kanno, K. (2005). Wing characteristics and flapping behavior of flying insects. Experimental Mechanics 45, 550-555.

Sudo, S., Tsuyuki, K. and Tani, J. (2000). Wing morphology of some insects. Jsme International Journal Series C-Mechanical Systems Machine Elements and Manufacturing 43, 895-900.

Wang, C. C. and Tang, D. J. (2009). Seafloor Roughness Measured by a Laser Line Scanner and a Conductivity Probe. Ieee Journal of Oceanic Engineering 34, 459-465.

Wilkin, P. J. and Williams, M. H. (1993). Comparison of the aerodynamic forces on a flying sphingid moth with those predicted by quasi-steady theory. Physiological Zoology 66, 1015-1044.

Wilkins, P. C. and Knowles, K. (2009). The leading-edge vortex and aerodynamics of insect-based flapping-wing micro air vehicles. Aeronautical Journal 113, 253-262.

Wood, R. J., Steltz, E. and Fearing, R. S. (2005). Optimal energy density piezoelectric bending actuators. Sensors and Actuators a-Physical 119, 476-488.

Wootton, R. J. (1990).The mechanical design of insect wings. Scientific American 263, 114-120.

Wootton, R. J. (1992). Functional-morphology of insect wings. Annual Review of Entomology 37, 113-140.

Yan, J., Wood, R.J., Avadhanula, S., Sitti, M., Fearing, R.S., Ieee, Ieee and Ieee. (2001). Towards flapping wing control for a micromechanical flying insect. In 2001 Ieee International Conference on Robotics and Automation, Vols I-Iv, Proceedings, pp. 3901-3908. New York: Ieee.

Zanker, J. M. and Gotz, K. G. (1990). The wing beat of drosophila-melanogaster .2. dynamics. Philosophical Transactions of the Royal Society of London Series B-Biological Sciences 327, 19-44.

Zeng, L. J., Matsumoto, H. and Kawachi, K. (1996). Simultaneous measurement of the shape and thickness of a dragonfly wing. Measurement Science and Technology 7, 1728-1732.

